# The scaffolding of individual variability in language processing by domain-general neural networks

**DOI:** 10.64898/2026.02.12.705531

**Authors:** Muge Ozker, Atsuko Takashima, Laura Giglio, Florian Hintz, Antje Meyer, Peter Hagoort

## Abstract

Language processing is supported by distributed neural systems. Yet most research examines these systems at the population-average level, obscuring how individual cognitive differences shape language-related brain activity. In this study, we combined comprehensive cognitive assessments and task-based fMRI in a large sample of healthy adults (N = 205) to examine how variability in linguistic knowledge, working memory, processing speed, and non-verbal reasoning influenced neural responses in four language tasks: lexical decision, picture naming, sentence comprehension, and sentence production. All tasks showed robust left-lateralized activation patterns and varied in the extent to which they engaged the classical perisylvian language network. At the level of individual differences, cognitive skills were not primarily associated with modulation of commonly activated language regions but largely associated with activity in domain-general systems outside these regions, particularly within the default mode, executive and attentional networks. Activation in these domain-general regions was predominantly negatively correlated with cognitive skills, indicating greater recruitment in individuals with lower cognitive skills. Overall, our findings demonstrate that individual cognitive profiles determine how distributed brain systems are dynamically engaged to scaffold language processing.

**Significance Statement:** How the human brain instantiates language across a diverse range of cognitive skills remains a fundamental mystery. Although language processing is typically attributed to a set of core language regions, we show that individual differences are not primarily reflected within these regions. Instead, variation in cognitive skills is largely associated with differential recruitment of domain-general brain networks, including default mode, executive control and attentional systems. Individuals with lower cognitive skill levels show greater engagement of these domain-general networks, consistent with a neural scaffolding mechanism that supports language processing. Together, these findings indicate that language is not a strictly modular function but is flexibly embedded within the brain’s broader, integrated neural architecture.

## Introduction

Producing even a single word engages a cascade of tightly coordinated processes, ranging from conceptualizing an intended message and retrieving the appropriate lexical item to assembling its phonological or gestural components into a precise motor plan (1–3). At the sentence level, these demands escalate: speakers must encode thematic roles (e.g., agent and patient), organize syntactic structure, and mark the communicative focus of the utterance (4).

Comprehension poses similarly complex challenges. Listeners must rapidly decode the incoming linguistic signal, integrate it with prior knowledge, and construct a coherent mental representation of the intended message in real time (5–8).

Successfully accomplishing these linguistic operations relies not only on access to domain-specific linguistic knowledge, i.e. vocabulary and grammatical knowledge (9, 10), but also on domain-general cognitive skills, including working memory, processing speed, and non-verbal reasoning. Working memory supports the maintenance and manipulation of transient conceptual and linguistic representations during both comprehension and production (11–15). For instance, it enables the retention of morphosyntactic structures and lexical items that are necessary for resolving long-distance dependencies (16, 17) and for integrating information across clause boundaries in complex sentences (18). Processing speed, defined as the efficiency with which cognitive operations are executed (19), constrains the ability to decode sensory input and plan linguistic output in real time. It is therefore critical for tasks requiring rapid temporal coordination, such as distinguishing phonemes in fast speech streams (20, 21) and retrieving and sequencing lexical candidates during fluent speech production (22). Non-verbal reasoning contributes to conceptual integration by supporting the construction of intended meaning beyond the literal words, the resolution of semantic ambiguities, the interpretation of metaphors, and the generation of discourse-level inferences (23, 24).

Importantly, domain-specific and domain-general skills vary substantially across individuals, raising the question of how such variability contributes to differences in language processing. Psycholinguistic research has consistently documented individual differences in language processing strategies and performance that are linked to cognitive skills. For example, working memory capacity has been associated with more efficient processing of syntactically complex sentences (25), while individual differences in processing speed have been linked to stronger linguistic prediction in visual-world paradigms (13, 26). In a similar vein, domain-general cognitive faculties extend to developmental contexts, where children with higher non-verbal reasoning skills have been shown to differentiate semantic and syntactic violations more rapidly (27). Together, these findings demonstrate that language performance is not determined by linguistic knowledge alone but is scaffolded by the flexible recruitment of domain-general cognitive resources that differ across individuals.

While there is mounting behavioral evidence for interindividual differences in language processing and growing interest in their neural correlates (28–32), neurobiological investigations of language have largely emphasized shared neural mechanisms across speakers and listeners (33–35). Consequently, individual variability has often been treated as noise rather than a source of meaningful insight into the neural mechanisms that support flexible language use (36). Traditional population-level and meta-analysis studies have consistently identified a canonical set of left-lateralized perisylvian regions implicated in language comprehension and production across both auditory and visual modalities. These regions include the inferior frontal gyrus (pars opercularis, triangularis, orbitalis), the posterior and middle superior temporal cortex, the middle temporal gyrus, anterior temporal lobes and the angular gyrus (37–44). A parallel line of research has used participant-specific language localizers, which accounts for interindividual variability in functional anatomy to improve the identification of language-selective regions that are often missed in conventional group analyses (45). However, a fundamental question regarding variability associated with broader cognitive skills remains unresolved: **How are individual differences in cognitive skills reflected in the neural processing of language?** We propose that at least two distinct neural mechanisms can be considered: One possibility is that individual differences may be expressed as variations in the degree of recruitment of the commonly activated domain-specific language regions. Alternatively, they may be reflected in the recruitment of additional domain-general regions that are engaged only by subsets of participants and therefore do not appear in the group-level activation maps.

Because linguistic and cognitive abilities develop in tandem and exhibit substantial interindividual variability, identifying the neural systems through which cognitive skills influence language processing can help explain why individuals with comparable language exposure still vary in proficiency and learning outcomes (46). Furthermore, establishing this relationship in neurotypical populations provides an important reference framework for interpreting compensatory neural recruitment in atypical development and clinical conditions. If domain-general systems contribute differentially to language processing across individuals with varying cognitive skills, similar mechanisms may contribute to resilience and recovery in conditions such as developmental language disorder, aphasia, and neurodegenerative disease.

In this study, we combined behavioral and neuroimaging approaches in a large sample of healthy participants (N = 205) for characterizing individual differences in language processing. Participants’ cognitive skills were assessed across four domains: domain-specific **linguistic knowledge,** and three domain-general skills, namely **processing speed, working memory**, and **non-verbal reasoning**. Additionally, during fMRI scanning, participants completed four language tasks: **auditory lexical decision, picture naming, sentence comprehension,** and **sentence production**, designed to tap different linguistic levels (words and sentences), sensory input modalities (auditory and visual) and task demands (comprehension and production). By relating interindividual variability in cognitive skills to variability in task-evoked brain activity, we aim to elucidate the cognitive and neural architecture that supports flexible language use across individuals.

## Results

### Individual Variability in Cognitive Skills

To characterize individual differences in cognitive skills, we used behavioral data previously collected from all participants as part of a comprehensive assessment of linguistic and domain-general cognitive abilities (47). From this battery, we derived latent factor scores capturing **linguistic knowledge, auditory processing speed, visual processing speed, and working memory** (**Table S1** provides an overview of the administered tests). **Non-verbal reasoning** was assessed separately using accuracy on the Raven’s Advanced Progressive Matrices test and was therefore represented by a single measure rather than a latent factor. Raven accuracy scores were standardized to place them on the same scale as the other cognitive measures. The distributions of all five measures across participants are shown in **Figure 1**. These individual-difference measures were subsequently included as covariates in the neuroimaging analyses.

**Figure 1.**
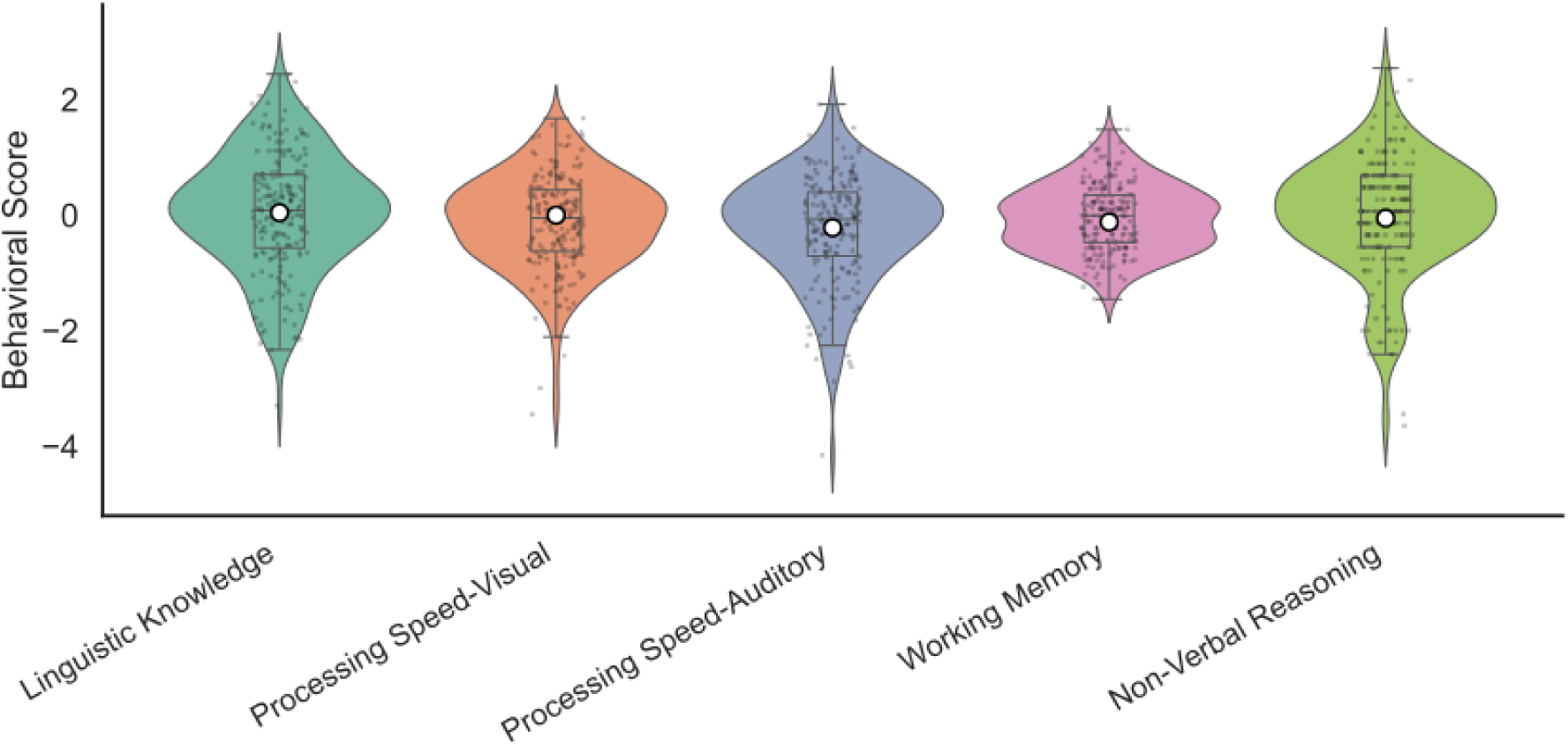
Distribution of behavioral scores across participants for different cognitive skills. Violin plots show the full distribution of individual scores for each cognitive skill, with overlaid boxplots indicating the median and interquartile range. Individual data points are jittered for visibility, and white markers with vertical lines represent the mean ± 95% confidence interval. Linguistic knowledge, processing speed, and working memory scores represent factor scores derived from multiple tests, whereas the non-verbal reasoning score corresponds to the z-scored accuracy on the Raven’s Advanced Progressive Matrices test.

### fMRI Results Across Language Tasks

We examined neural responses during four language tasks spanning word- and sentence-level comprehension and production: **auditory lexical decision**, **picture naming**, **auditory sentence comprehension**, and **sentence production**. Participants made active responses in the scanner, either through button presses (lexical decision and sentence comprehension) or overt speech (picture naming and sentence production). These responses were analyzed to confirm participant engagement and task compliance. Accuracy scores were consistently high across all tasks and conditions (> 90%, **Table S2**), indicating that participants were attentive and reliably followed the task instructions.

Next, we conducted two complementary group-level analyses to examine neural activation patterns during the four language tasks: one aimed at identifying **common activation across participants**, and the other aimed at identifying neural responses associated with **individual differences** in cognitive skills.

We examined a targeted contrast of interest within each language task to isolate specific linguistic processes. For the lexical decision task, we focused on the ***Word > Quilt*** contrast. The Quilt stimuli retained the low-level auditory features of real words but lacked recognizable phonotactic or lexical structure, allowing this contrast to control for basic auditory input while isolating brain regions involved in phonological and lexical processing. In the picture naming task, we used the ***Object > Nameless Object*** contrast. Because both conditions required spoken responses, this comparison controlled for general speech-related motor activity and highlighted regions specifically engaged in lexical access and semantic processing. For the sentence comprehension task, we examined the ***Sentence > Wordlist*** contrast. Only the Sentence condition contained coherent syntactic and semantic structure, making this contrast sensitive to higher-level integration processes. Finally, in the sentence production task, we analyzed the ***Sentence > Phrase*** contrast. As both conditions involved speech production, this contrast isolated neural regions involved in generating full, syntactically structured and semantically integrated utterances, beyond the demands of producing isolated phrases.

In addition, for the sentence comprehension and sentence production tasks, we examined how brain activity was modulated by **syntactic complexity** using parametric analyses. In sentence comprehension, we modeled activation as a function of left-branching unification demands to identify regions sensitive to structure building. In sentence production, we used a linear modulator to capture increasing syntactic complexity across conditions (simple phrases < intransitive < active < passive sentences), revealing regions that support the construction of increasingly complex syntactic representations. However, because sentence length in this task increased systematically across conditions, the resulting effects reflect the combined influence of syntactic complexity and increasing utterance length.

### Common activations across individuals

Focusing first on common activation patterns, a whole-brain comparison of the four language tasks revealed both overlapping and task-specific neural activation profiles. For each task contrast, group-level statistical T-maps were thresholded to identify significant positive activations (*T* > 3.13, *p* < 0.001 uncorrected). The extent of activation, measured by the number of suprathreshold voxels, varied across tasks: picture naming elicited the broadest activation (47,417 voxels), followed by sentence comprehension (45,401), lexical decision (32,595), and sentence production (28,671) (**Figure 2**).

**Figure 2.**
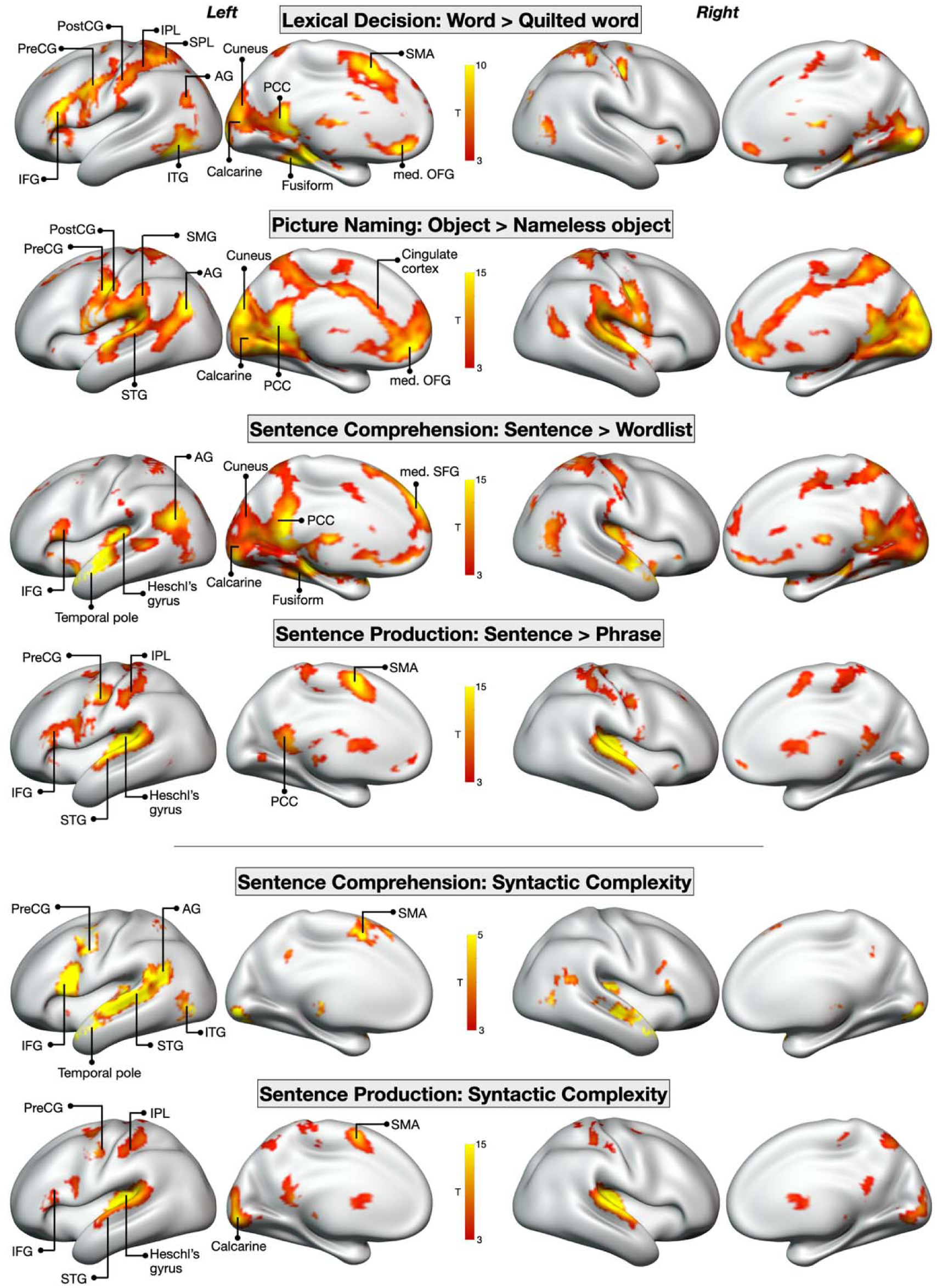
fMRI group-level activation patterns for different language tasks. Cortical surface renderings (HCP S1200 32k_fs_LR inflated surface) showing group-level common activation patterns across contrasts and parametric modulators for each language task. Pre/PostCG: Precentral/Postcentral gyrus, IFG: Inferior frontal gyrus, ITG: Inferior temporal gyrus, AG: Angular gyrus, med. OFG: medial orbital frontal gyrus, SMA: Supplementary motor area, SMG: Supramarginal gyrus, STG: Superior temporal gyrus, med. SFG: medial superior frontal gyrus, IPL: Inferior parietal lobule, SPL: Superior parietal lobule, PCC: Posterior cingulate cortex

To compare the activation patterns for each language task between the left and right hemispheres, we computed a laterality index (LI) for each task with positive LI values indicating left-lateralization (see *Laterality Index Calculation*). All four language tasks demonstrated significant left-hemisphere asymmetry in activation, consistent with the broadly observed left-lateralization of language processing in the brain (48, 49). However, the degree of lateralization varied by task, indicating that different tasks modulated the relative contributions of the two hemispheres. The sentence comprehension tasks showed the strongest left-lateralization (LI = 0.34, t = 18.32, p < 0.001, Cohen’s d = 1.32) followed by lexical decision (LI = 0.24, t = 8.66, p < 0.001, Cohen’s d = 0.62), picture naming (LI was 0.22, t = 11.98, p < 0.001, Cohen’s d = 0.87) and sentence production (LI = 0.16, t = 11.52, p < 0.001, Cohen’s d = 0.83).

To identify common activation clusters for each task contrast, we applied a voxel-wise threshold of *p* < 0.001, followed by cluster-level Family-Wise Error (FWE) correction to control for multiple comparisons (**Figure 2, Table S3**). The **lexical decision** task elicited large activation clusters centered in the posterior portion of the left inferior temporal gyrus (ITG), extending into the posterior middle temporal gyrus (MTG), fusiform gyrus, posterior cingulate cortex, cuneus and calcarine. Strong activation was also observed in the inferior and superior parietal lobules (IPL and SPL) and inferior frontal gyrus (IFG), centered in pars orbitalis (50). This pattern aligns with findings from previous meta-analyses of speech perception comparing spoken words and pseudowords, which identified a similar network supporting phonological processing and lexical-semantic access to extract meaning from spoken words (51, 52).

The **sentence comprehension** task also elicited activation in the posterior ITG, extending into the fusiform gyrus and medial visual regions. However, unlike the lexical decision task, which primarily engages word-level comprehension, we also observed robust bilateral activation of the temporal poles (anterior temporal lobe, ATL). This pattern aligns with previous studies comparing sentence comprehension to unstructured word lists, highlighting the ATL’s role in processing syntactic structure and combinatorial semantics (53–56). Additionally, we observed activation in the medial prefrontal and posterior cingulate cortices during sentence comprehension. These regions, typically implicated in episodic memory retrieval (Rugg & Vilberg, 2013), have also been reported to be recruited during tasks that require encoding sentence content into memory (59). This is consistent with our task design, in which participants answered comprehension or memory probe questions on selected trials.

Tasks that required overt articulation, namely **picture naming** and **sentence production**, engaged bilateral frontal motor and sensorimotor regions in the precentral and postcentral gyrus, as well as the cerebellum, consistent with the demands of motor planning and execution. Notably, both tasks also elicited extensive activation in the right auditory cortex, encompassing Heschl’s gyrus and extending into the superior temporal gyrus (STG), indicating a prominent role for auditory feedback monitoring during speech production. Sentence production further recruited the IFG, and supplementary motor area (SMA), supporting its involvement in motor sequence control and coordination (60). Meanwhile, picture naming elicited significant activation in occipital regions, including the calcarine cortex, cuneus, lingual gyrus, and occipital gyrus, reflecting the visual processing demands of the task necessary for object recognition.

### Activations that reflect individual differences

Next, we identified neural responses associated with individual differences in cognitive skills, i.e. linguistic knowledge, processing speed, working memory, and non-verbal reasoning. To do this, we conducted group-level analyses using participants’ behavioral scores (**Figure 1**) in these four cognitive domains as covariates. Significant activation clusters were defined using a voxel-wise threshold of *p* < 0.001, followed by cluster-level FWE correction. This analysis revealed several clusters where activation levels varied systematically with cognitive skill levels (Clusters 1-25 in **Table S4** and **Figure 3**). These ‘individual variability clusters’ were distributed across both hemispheres (17 left, 8 right). The largest number of clusters was identified for the sentence comprehension and sentence production tasks, accounting for 18 of the 25 clusters. Across cognitive domains, clusters were most frequently associated with linguistic knowledge (11 of 25 clusters) and least frequently associated with working memory (3 of 25 clusters). Sixteen of the 25 clusters showed negative correlations with cognitive skills, indicating that participants with lower cognitive skills exhibited higher activation in these regions.

**Figure 3.**
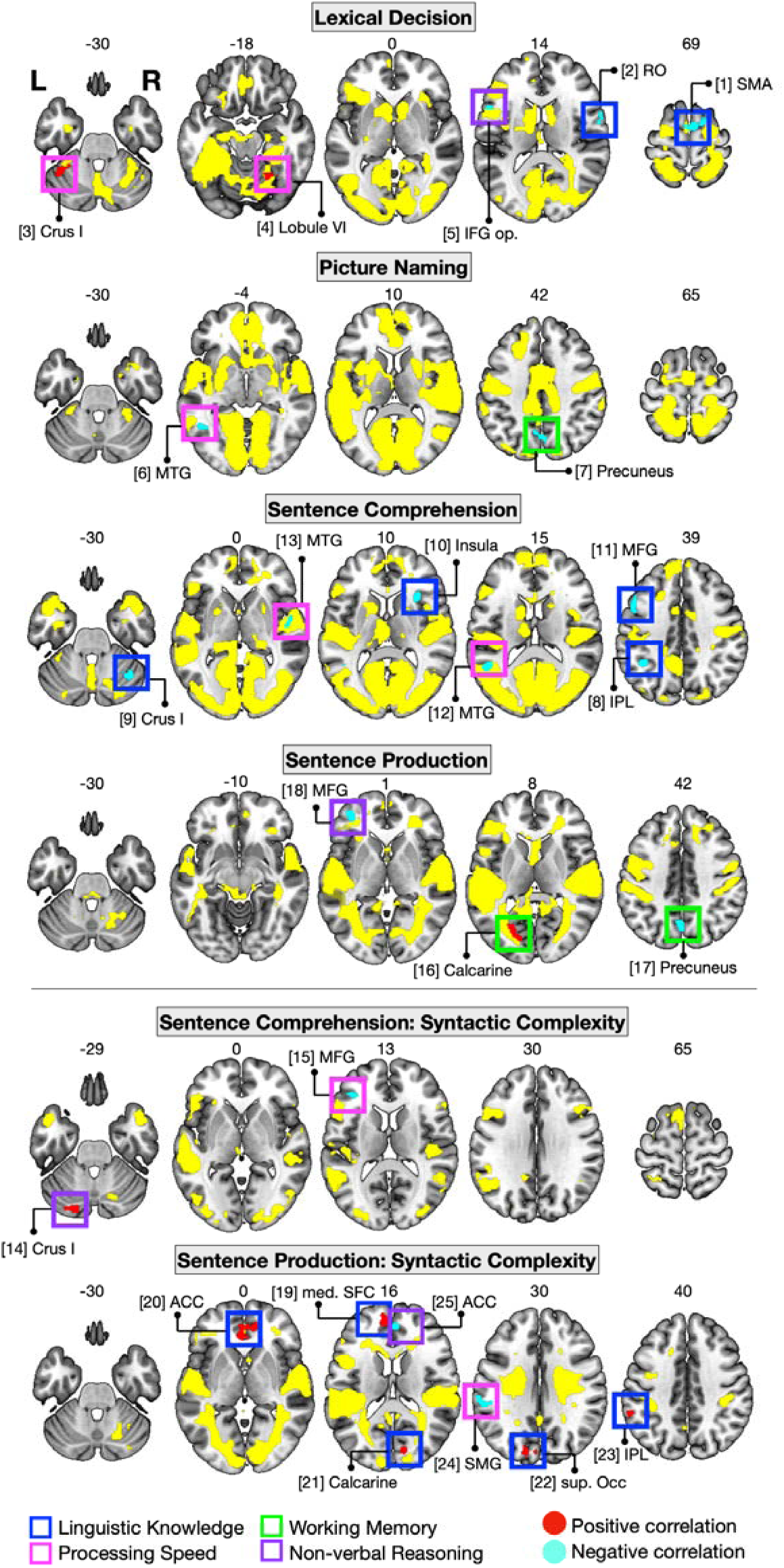
fMRI activation patterns for common effects and individual differences. Yellow indicates group-level common activation patterns across contrasts and parametric modulators for each language task. Five axial slices are displayed for each task, with slice numbers indicated on top (MNI 152). Red and blue clusters denote regions where activation correlates with behavioral scores (Red = positive correlation; Blue = negative correlation). These clusters are highlighted by colored square frames, where the color of the frame indicates the specific cognitive skill associated with the effect. Cluster numbers 1-25 correspond to those in **Table S4**. SMA: Supplementary motor area, RO: Rolandic Operculum, IFG op.: Inferior frontal gyrus opercularis, MTG: Middle temporal gyrus, STG: Superior temporal gyrus, MFG: Middle frontal gyrus, SMG: Supramarginal gyrus, AG: Angular gyrus, IPL: Inferior parietal lobule, med. SFC: Medial superior frontal cortex, ACC: Anterior cingulate cortex, sup. Occ: superior occipital gyrus

We then examined the anatomical locations of these clusters relative to the group-level common activation maps to determine whether they were colocalized with shared activation regions or located outside them. To quantify this relationship, we calculated the percentage of spatial overlap between each cluster and the common activation map of the corresponding language task. Although some overlap was observed, only 3 of the 25 covariate clusters showed more than 50% spatial overlap with the commonly activated regions. Six clusters were adjacent to commonly activated regions but showed only minimal overlap (2-26%), whereas 16 clusters showed no overlap. Thus, the majority of regions associated with individual differences in cognitive skills showed little or no spatial overlap with the group-level activation patterns (**Table S4, Figure 3**). The greatest degree of overlap was observed for the lexical decision and sentence comprehension tasks, where clusters overlapped with regions including the IFG, STG, MTG, SMA, and cerebellum. For sentence production, only limited overlap with the calcarine cortex was observed, while all other clusters across tasks fell entirely outside the shared activation regions. These results suggest that individual differences in language processing are largely associated with the recruitment of brain regions that are distinct from, or only partially overlapping with, regions commonly engaged across participants.

### Brain networks underlying individual variability in language processing

To gain insight into the large-scale brain systems that contribute to individual variability in language processing, we next examined how the previously identified ‘individual variability clusters’, in which activation during language tasks covaried with cognitive skills, mapped onto established functional networks. Specifically, we quantified the spatial overlap between each cluster (Clusters 1-25 in **Table S4** and **Figure 3**) and the seven canonical Yeo networks (61): Visual Network (VN), Somatomotor Network (SMN), Dorsal Attention Network (DAN), Ventral Attention Network (VAN), Limbic Network (LN), Frontoparietal Network (FPN), and Default Mode Network (DMN). The Yeo networks provide a well-established and widely used functional parcellation of the cortex, offering a principled framework to characterize how individual variability relates to large-scale neurocognitive systems.

Across tasks, eight clusters overlapped with the DMN, with more than 50% of each cluster falling within the network, highlighting DMN’s central role in supporting the high-level cognitive operations that contribute to language processing. Four clusters similarly showed more than 50% overlap with the VAN and four with the FPN, pointing to the role of attentional and executive control. The DAN, SMN and VN were also involved but to a lesser extent, whereas none of the clusters showed overlap with the LN (**Figure 4**).

**Figure 4.**
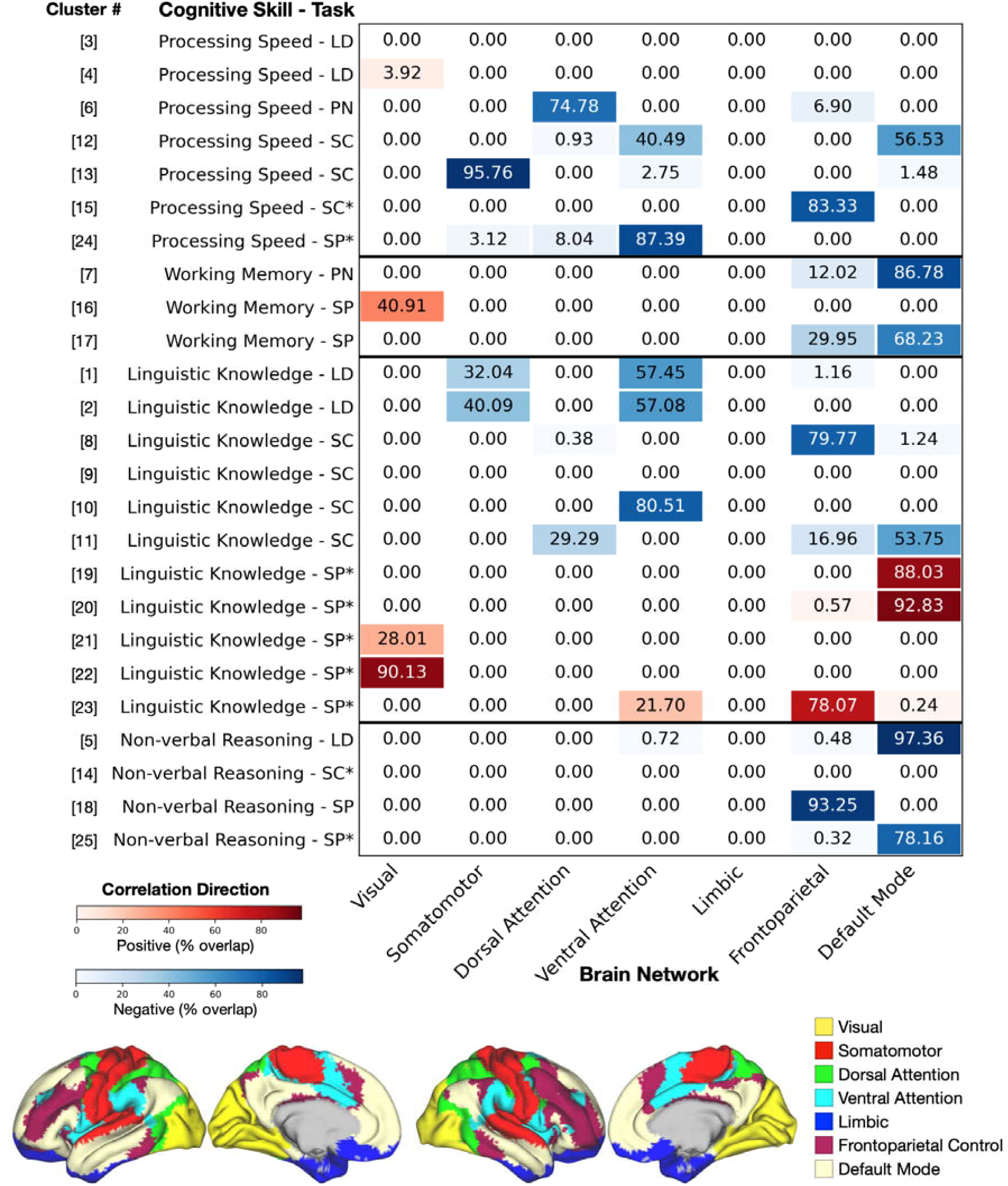
Spatial overlap of fMRI clusters reflecting individual variability with the seven canonical Yeo networks. Cluster numbers correspond to those in **Table S4** and Figure 3. Red and blue denote positive and negative correlations, respectively, between cluster activation and participants’ behavioral scores across cognitive skills, while color intensity indicates the degree of overlap between each cluster and the network. LD: Lexical Decision, PN: Picture Naming, SC: Sentence Comprehension, SC*: Sentence Comprehension (Syntactic Complexity), SP: Sentence Production, SP*: Sentence Production (Syntactic Complexity)

Distinct profiles emerged when examining network overlap patterns across the four cognitive domains. Clusters associated with **linguistic knowledge** showed a broadly distributed profile, spanning diverse anatomical regions and overlapping with most functional networks to varying degrees (**Table S4, Figures 3-4**). Similarly, clusters associated with **processing speed** were localized to a range of regions including the cerebellum, MTG, STG, MFG, and SMG, and distributed across multiple networks. In contrast, the remaining two cognitive skills showed more selective network profiles. Clusters associated with **working memory** emerged exclusively during overt speech production tasks and were primarily located in the left precuneus and calcarine cortex, with overlap mainly involving the DMN and FPN. Finally, clusters associated with **non-verbal reasoning** also exhibited a focused profile, localizing to the medial prefrontal/anterior cingulate cortex and the IFG (opercularis), with overlap restricted to the DMN and FPN.

## Discussion

Our study examined how individual differences in cognitive skills are reflected in neural activity during language processing. Across language tasks, group-level activation patterns showed a consistent left-lateralized organization, yet the involvement of classical perisylvian language regions varied. Notably, clusters associated with individual differences in cognitive skills were largely located outside the commonly activated regions, most prominently involving the default-mode, frontoparietal and ventral attention networks. The majority of these clusters showed negative correlations with cognitive skills, indicating that individuals with lower cognitive skills exhibited stronger engagement of these domain-general regions, suggesting that broader neural resources may be recruited to support language processing.

Although all four language tasks showed robust left-lateralized activation patterns, they differed in the extent to which they engaged the classical perisylvian language areas. For example, the lexical decision task activated the left IFG but did not elicit significant activation in the posterior STG/STS. Conversely, some tasks recruited additional regions beyond the canonical language areas; for instance, picture naming and sentence comprehension elicited extensive activation in the medial occipital cortex. These differences in activation profiles likely reflect variation in task demands and the specific contrasts used in our study, as well as the inherent challenge of isolating linguistic processes from the sensorimotor and cognitive processes that accompany language tasks.

The degree of left lateralization also varied across tasks. Sentence comprehension, for example, elicited slightly stronger left-lateralized activation (LI = 0.34) than word-level processing in the lexical decision task (LI = 0.24), consistent with models proposing greater left-hemisphere dominance for connected speech compared with single-word processing (62, 63). Right-hemisphere regions, including Heschl’s gyrus and the superior temporal gyrus (STG), as well as the cerebellum, were also strongly engaged during overt speech production, underscoring the contribution of bilateral systems to sensorimotor coordination and speech monitoring (64–66).

Although both sentence production and picture naming involved visually presented stimuli, picture naming additionally elicited stronger right-hemisphere visual activation, possibly reflecting greater object-recognition demands of the task (67). In contrast to the auditory comprehension tasks, production tasks exhibited a different pattern of lateralization: word-level processing in the picture naming task showed stronger left lateralization (LI = 0.22), whereas sentence production recruited relatively more bilateral networks (LI = 0.16). Together, these findings support the view that hemispheric asymmetry in language is dynamic and shaped by input modality (visual vs. auditory), task demands (production vs. comprehension), and linguistic level (word vs. sentence processing)(43, 68).

Importantly, individual variability in language processing did not entirely manifest within these shared patterns of task-related activation. Instead, correlations between brain activity and cognitive skills revealed that variability across individuals was largely expressed in regions showing little overlap with, or extending beyond, the group-level activation maps. Furthermore, the majority of activation clusters associated with cognitive skills showed negative correlations with cognitive skills. The predominance of these negative associations suggests that individuals with lower cognitive skill levels may engage broader neural resources more strongly during language processing, whereas higher-skilled individuals achieve comparable performance with reduced auxiliary engagement.

Characterizing these effects within the framework of large-scale functional brain networks provided additional insight into the neural systems through which individual variability emerges. Clusters associated with **linguistic knowledge** spanned multiple functional networks and, with the exception of the sentence production task, showed a consistent pattern of reduced neural engagement with increasing linguistic knowledge. Positive associations between cognitive skills and neural activity were uncommon and were observed exclusively for linguistic knowledge during the sentence production task. During the production of complex sentences, individuals with higher linguistic knowledge showed greater activation within the visual network (VN), frontoparietal network (FPN), and default mode network (DMN). In contrast, during lexical decision tasks, higher linguistic knowledge was associated with reduced activation in frontal motor regions overlapping with the ventral attention network (VAN) and sensorimotor network (SMN), implying that individuals with more extensive linguistic knowledge recognize words more automatically, requiring less attentional or motor simulation effort. Conversely, individuals with lower linguistic knowledge may recruit these regions more strongly, possibly reflecting compensatory strategies such as increased attentional effort or subvocal rehearsal.

Clusters associated with **processing speed** also spanned a broad range of networks, including regions involved in sensorimotor coordination, visuospatial processing, and multisensory integration, with overlap across the SMN, VAN, DAN, FPN and default mode network (DMN). This widespread network involvement suggests that rapid language processing depends on the coordinated interaction between perceptual systems and domain-general control mechanisms that support flexible integration across modalities.

In contrast to the broad network involvement observed for linguistic knowledge and processing speed, **working memory** and **non-verbal reasoning** showed more selective network engagement, primarily involving the FPN and DMN. Working memory skills were particularly linked to speech production tasks, with clusters located in the precuneus, a DMN hub associated with internally oriented cognition, memory integration, and construction of mental representations (57, 69). Because both production tasks in our study relied on visually presented stimuli, increased precuneus activation in individuals with lower working memory capacity may reflect a greater reliance on DMN-mediated processes to transform visual input into mental representations that support verbal planning and short-term retention (70). Similarly, non-verbal reasoning was negatively associated with FPN and DMN engagement, particularly in the medial prefrontal and anterior cingulate cortices, regions implicated in abstraction, strategic reasoning, and goal maintenance (71). These findings indicate that low-level reasoning skills modulate engagement with linguistic input, even in the absence of explicit problem-solving demands when the task is primarily linguistic.

Across cognitive domains, the DMN, FPN, and attention networks emerged as the networks most consistently associated with individual differences. Lower working memory and non-verbal reasoning skills were associated with greater activation within the DMN and FPN, whereas lower linguistic knowledge and processing speed were associated with greater engagement of attentional networks. The DMN is thought to play a central role in language processing by integrating multimodal information into coherent, internally generated representations of meaning. During language tasks, it supports the construction of conceptual and semantic models that link words to prior knowledge and context (70, 72, 73). Reduced DMN activation observed in individuals with higher working memory and non-verbal reasoning skills, suggests that they can access and manipulate linguistic representations with less reliance on broad, integrative mechanisms. In contrast, individuals with lower skills may depend more heavily on DMN-mediated processes to maintain and bind information. Similarly, greater VAN and DAN activation in individuals with lower linguistic knowledge or slower processing speed may reflect increased reliance on attentional control mechanisms. Attentional networks have long been implicated in language processing, supporting the selective allocation of attention to semantic categories (74) and to specific acoustic features such as pitch, intonation, lexical tone, and prosodic focus (75–78). Greater engagement of these networks in lower-skilled individuals likely reflects the need for sustained attentional support to extract linguistic cues and integrate information over time, processes that higher-skilled individuals perform more automatically and efficiently.

Overall, by combining cognitive profiling with task-based fMRI across multiple language tasks, we show that variability associated with cognitive skills is expressed predominantly in distributed brain regions that extend beyond the shared task-related activation patterns. These regions are largely associated with domain-general networks, particularly the default mode, executive, and attentional networks, suggesting that individual differences in cognitive skills are reflected primarily in the recruitment of these networks rather than in modulations of the commonly activated language regions. While our group-level activation maps are consistent with established models emphasizing the central role of core language regions (79–81), the present findings suggest that individual differences in language processing are also associated with variation in the engagement of broader cognitive systems. Together, these findings support the view that understanding individual variability in the neural basis of language requires considering both the shared language network and the contribution of domain-general cognitive networks, providing a framework for interpreting interindividual differences in functional neuroimaging studies of language.

Nevertheless, several limitations should be considered when interpreting the conclusions that can be drawn from the present findings. First, participants performed the language tasks with high accuracy, which may have reduced the variability in task performance and limited our ability to capture neural correlates associated with lower levels of language proficiency or performance. Previous research in both developmental and aging populations suggests that individual differences in cognitive abilities such as working memory, executive control, and language skills become increasingly important as processing demands increase. For example, studies of sentence comprehension in children have shown that developmental differences in language and cognitive abilities modulate neural responses, particularly as sentence complexity increases (82, 83). Similarly, studies in healthy older adults have demonstrated that individual differences in working memory, hearing, and cognitive control predict variability in neural responses during language comprehension (84, 85). Although we observed reliable associations between cognitive abilities and neural responses despite near-ceiling performance, future work should determine whether stronger or additional effects would emerge in more demanding language paradigms or in younger and older populations, where cognitive constraints play a larger role in shaping language processing.

Another important direction for future research will be to examine whether the associations observed here generalize to more naturalistic language contexts. The present study focused on a set of well-established, yet highly controlled tasks that allow specific linguistic processes to be isolated and compared across individuals. However, these tasks place relatively modest demands on the integration of information over extended timescales. In contrast, naturalistic language comprehension, such as narrative listening, requires the continuous integration of information across extended timescales and engages a broader set of cognitive processes, including working memory, inference, attention, and prior knowledge, that support the construction of coherent situation models (21, 86, 87). Accordingly, individual differences in cognitive skills may play an even larger role in shaping neural responses under these more ecologically valid conditions. We are currently analyzing the narrative comprehension data acquired as part of the present project (88) and we anticipate that these analyses will complement the present findings by revealing how cognitive variability influences neural processing during continuous, real-world language comprehension.

## Materials and Methods

### Participants

A total of 205 healthy young adults, who were native Dutch speakers, participated in the study (age range: 18-30; mean age = 23 years; 149 females; 28 left-handed; 3 ambidextrous). Inclusion criteria were as follows: (1) fluent Dutch speakers; (2) normal hearing and normal vision or corrected-to-normal vision; (3) no history of brain operations or epilepsy; (4) fulfill the safety criteria to go into the MRI scanner. Several participants were excluded due to anatomical anomalies, poor task performance, or missing behavioral data, resulting in the following final sample sizes: Lexical decision (n = 193), picture naming (n = 190), sentence comprehension (n = 192), and sentence production (n = 192). The study was approved by the local ethical committee of the Faculty of Social Sciences at Radboud University (ECSS/ECSW; CMO2014/288), and written informed consent was obtained from all participants in accordance with the Declaration of Helsinki.

### Behavioral Testing and Cognitive Skill Assessment

All 205 participants were recruited from a larger pool of 748 individuals who had previously completed a comprehensive behavioral study designed to assess interindividual differences in linguistic knowledge, linguistic processing skills, and domain-general cognitive skills using an extensive test battery designed by Hintz et al. (47). Six participants completed the behavioral battery on-site (two-hour sessions on two separate days), while the remaining 199 participants completed the same battery online (four one-hour sessions on four separate days). For the present study, we used latent factor scores derived from tests of linguistic knowledge, auditory and visual processing speed, and working memory, following the procedure described in Hintz et al. (47). Non-verbal reasoning was assessed using accuracy on the Raven’s Advanced Progressive Matrices test and was therefore represented by a single measure rather than a latent factor (**Table S1**). Tests targeting linguistic processing abilities (i.e., word- and sentence-level comprehension and production) were not included in the present analyses and were distinct from the language tasks performed during fMRI. provides an overview of the behavioral tests included in the battery.

### fMRI Testing

All participants attended a single three-hour laboratory session, divided into two halves. In the first half, they completed the **lexical decision** and **picture naming** tasks, and in the second half, they performed the **sentence comprehension** and **sentence production** tasks (*see SI Appendix for a detailed description of experimental procedures*). Participants also listened passively to two 4-minute stories, one at the end of each session half, but data from the stories were not analyzed in this study.

### Lexical Decision Task

#### Stimuli

The stimuli consisted of 80 existing Dutch nouns, 80 pseudowords and 20 quilts. Recordings of all the existing words and pseudowords were produced by one Dutch female speaker. Quilts were created by dividing real word recordings into segments of 30 ms in length, which were then concatenated in a semi-random order, which essentially disrupted sound properties of audio stimuli on longer timescales, while approximately preserving them on shorter timescales. This means that quilted stimuli maintained their low-level auditory features (i.e., their speech-specific spectrotemporal structure), without any recognizable phonotactic or lexical features.

### Procedure

Participants performed the auditory lexical decision task in the scanner with an event-related design, consisting of two runs: 40 word, 40 pseudoword and 10 quilt trials per run. There was a short break of 30 s between the two runs while the scanner continued collecting the data. The trial structure was as follows: participants were presented with a red fixation cross in the middle of the screen for 250 ms, indicating the beginning of a trial, followed by the presentation of the auditory stimulus (word, pseudoword, or quilt). The trial length including the auditory input was 2750 ms. Participants were instructed to press the “yes” button with their left index finger when hearing an existing word and press the “no” button with their left middle finger when hearing non-existing words or quilts. After the response was given, the red fixation cross turned black. The inter-trial interval was jittered between 1.5 s to 6 s in steps of 1.5 s, 2.1 s on average. The whole task including the 30 s pause lasted approximately 13 min (**Figure S1**). The order of trials was pseudo-randomized, but the same order was used for all participants. “OptSeq” software was used to randomize and optimize the order of trials and the inter-trial interval (https://surfer.nmr.mgh.harvard.edu/optseq/) (89).

### Analysis

At the individual subject level, three regressors of interest (trials of words, pseudowords, and quilts) that were responded to correctly, along with a regressor for incorrectly responded trials, were modeled using the Hemodynamic Response Function (HRF). A fifth regressor accounted for the 30 s pause halfway through the task. Furthermore, motion-related nuisance regressors were added to the model. For the group level analysis, we focused on the **Word > Quilt** contrast, as these two conditions were acoustically similar. At the group level, we investigated how individual differences in cognitive skills influenced activation patterns. A random-effects analysis was conducted, using each participant’s contrast images as random factors. Four behavioral factor scores were included as covariates of interest: linguistic knowledge, auditory processing speed, working memory, and non-verbal reasoning. Gender and handedness were also included as covariates of no interest.

### Picture Naming Task

#### Stimuli

The stimuli consisted of 80 pictures of common objects, 20 nameless objects and 20 scrambled pictures. Scrambled pictures were generated by dividing the intact pictures into a 20*20 grid of small squares (each 15*15 pixels), and by randomly rearranging the positions of these small squares. The resulting scrambled pictures were thus similar to the original pictures in terms of low-level visual features and complexity, but without any recognizable parts or objects. Twenty nameless objects were selected from other studies that used such objects in their studies (90, 91). This condition required the visual processing of picture stimuli and generating a verbal response (“Niks”) without engaging in accessing, retrieving, or preparing lexical representations for production.

### Procedure

The run consisted of two halves with each half having 40 objects, 10 nameless objects, and 10 scrambled pictures with a 30 sec pause in between while the scanner continued to acquire volumes. Participants were instructed to name the object using one word, without articles, in singular form, and not using diminutives. If participants saw a nameless object, they were instructed to say “Niks” (“nothing” in Dutch). For the scrambled pictures, participants were instructed to stay silent. The trial structure was as follows: a black cross appeared in the middle of the screen for 150 ms, indicating the beginning of a trial, followed by the presentation of the visual stimulus (object, nameless object, or scrambled picture). The picture presentation was fixed to 2850 ms. The inter-trial interval was jittered between 1.5 s to 6 s in steps of 1.5 s, 2.4 s on average (**Figure S2**).

### Analysis

As regressors of interest, correct trials for object, nameless object, and scrambled picture conditions, and incorrect trials of all conditions pooled together, and a fifth regressor to account for the pause halfway through the task were included. Furthermore, motion-related nuisance regressors were added to the model. For the group level analysis, we focused on the **Object > Nameless Object** contrast, as participants responded by speaking in both conditions. At the group level, a random-effects analysis was conducted using the same approach and covariates as described for the Lexical Decision task.

### Sentence Comprehension Task

#### Stimuli

80 sentences were selected from a published data set Mother Of Unification Studies (MOUS) created by Schoffelen et al. (92) and were used in a study by Uddén et al. (93). All sentences varied between 9 and 15 words in length, and they were relatively unconstrained in terms of syntactical structure. For the word list condition, 140 words were randomly selected from the list of sentences. These words were distributed amongst 20-word lists with seven words in each list. The eighty spoken sentences had an average duration of 4063 ms (range 2588 - 5147) and 20-word lists had an average duration of 5443 ms (range 4609 – 6277 ms). In 20% of the cases (16 trials for sentence condition, and 4 trials for word list condition), a question followed on the screen, which focused on the content of the sentence, or whether the word was mentioned in the preceding sentence / list. (e.g. “Was a profession mentioned?”, “Was the word ‘barkeeper’ mentioned?”). All sentences were additionally characterized in terms of syntactic complexity, as well as word count. The syntactic complexity of each sentence was assessed using a left-branching complexity measure from a dependency structure (93, 94).

### Procedure

On each trial, participants first saw a fixation cross in the middle of the screen for 500 ms, indicating the start of a trial. This was followed by the presentation of a spoken sentence on task trials, or a spoken random word lists on the baseline trials. Participants were instructed to listen to the sentences carefully and to keep their attention on the task, twenty percent of the stimuli were followed by a visually presented comprehension question on sentence trials or a memory probe question on word list trials. Participants were instructed to make a “yes” or “no” judgment as quickly and accurately as possible by pressing a button associated with left index (yes) or left middle finger (no), while the question remained visible on the screen for 6 s. The inter-trial intervals were jittered between 1.5 s to 6 s, 2.4 s on average. The order of trials was pseudo-random and the same for each participant. Halfway through the run, there was a 30 s pause where the participants were instructed to stay still in the scanner. The experiment lasted approximately 17.6 min (**Figure S3**).

### Analysis

Two regressors of interest (sentence condition, wordlist condition) and two control regressors (question and pause) were modelled together with the motion-related nuisance regressors. All experimental conditions were modeled using the onset times and durations of the sentence, word list, and question trials, convolved with the canonical hemodynamic response function (HRF). The pause regressor was modelled using the onset of the pause with a duration of 30 s. As a measure of syntactic complexity, left-branching (LB) unification scores were included as a parametric modulator for the sentence condition, after controlling for sentence length (word count) (94). For the group-level analysis, we focused on the **Sentence > Word List** contrast and the effect of LB syntactic complexity. The random-effects analyses were conducted using the same approach and covariates as described for the Lexical Decision task.

### Sentence Production Task

#### Stimuli

The stimuli were adapted from the set of photographs used in the studies by Menenti et al. (95, 96). The stimulus set included 30 images depicting a single person shaded in yellow performing an action, 30 images showing two people (a man and a woman) engaged in an action with one shaded in yellow and the other in blue, and 30 additional images featuring the same pair performing actions with the color assignments swapped between the man and the woman. In addition, 30 intransitive, 30 active and 30 passive sentences, and two phrases spoken by a native Dutch young male were used. In total there were 30 active trials, 30 passive trials, 30 intransitive trials and 24 phrase trials (baseline) (**Figure S4A**).

### Procedure

In order to guide the active vs. passive voice production in the experimental trials (transitive events), we implemented a combination of structural priming and the so-called stoplight paradigm, as described by Menenti et al. (96). That is, the pictures were preceded by an audio recording of a sentence that featured the same syntactic structure as the target sentence, using a different verb to prime the participant. The participants were not instructed to copy the sentence structure of the audio material but were instructed to produce one sentence per picture using the verb on the screen and mentioning the yellow person first. This coerced the production of active (e.g. The woman pushed the man – when the woman was yellow) and passive (e.g. The man was pushed by the woman – when the man was yellow) sentences. The actor on the intransitive condition (e.g. The man yawns) was always shaded in yellow. For the baseline phrase condition, a color-word, either “geel (yellow)” or “blauw (blue)” was presented instead of a verb. The participants were instructed to name the person in that color (e.g., “de man (the man)” or “de vrouw (the woman)”). The preceding audio for this phrase condition was either “de man of de vrouw (the man or the woman)” or “de vrouw of de man (the woman or the man)”.

The trial began with a fixation cross for 200 ms in the middle of the screen. Then, while the fixation cross remained in view, they heard the priming sentence (between 1000 and 3000 ms in duration), which indicated the structure of the target sentence (active, passive, or intransitive). Then, the target verb or the color word was presented visually for 500 ms, followed by another fixation cross lasting 200 ms. After that, the target picture was presented, and participants had to describe the event as quickly and accurately as possible, using the color coding in the picture.

The inter-trial interval was jittered between 1.5 seconds and 9 seconds, with an average of 2.7 seconds. The order of trials was pseudo-random and the same for each participant, such there was always an intransitive condition or a phrase condition in between an active or a passive condition except for six instances where the passive condition was directly followed by an active condition (4 times) or a passive condition (2 times). The experiment consisted of two halves with a 30 s break in between. The conditions were divided equally between the two halves and the whole experiment lasted for approximately 20 min (**Figure S4B**).

### Analysis

At the single subject level, we modelled three levels of syntactic complexity (intransitive, active, and passive) and one baseline condition (phrase) for all correctly produced trials, and three control regressors, one accounting for the prime sentence period, one accounting for all incorrect trials, and one for the pause block. For each trial, speech onset and offset times were extracted from the participants’ in-scanner voice recordings. Experimental regressors were modelled using trial durations from picture onset to speech offset. In addition, speech onset was included as a parametric modulator for each experimental condition. In this model, we extracted contrast images for the effect of sentence production (including intransitive, active and passive conditions) relative to the baseline phrase production condition. In a second model, we modelled four regressors, one for correct trials, one for incorrect trials, one for the prime sentence period and one for the pause block. Correct trials were modelled using trial durations from picture onset to speech offset, and speech onset was included as a parametric modulator. Furthermore, we incorporated a parametric modulator, which assigned weights to the syntactic complexity in a linear fashion (-1.5 for phrase condition, -0.5 for intransitive, 0.5 for active, and 1.5 for passive sentence conditions) to the correct trial condition. Both models included motion related nuisance regressors like the other three tasks. For the group level analysis, we focused on the **Sentence > Phrase** contrast from the first model, and the effect of syntactic complexity from the parametric modulator in the second model. Random-effects analyses were conducted using the same approach and covariates as described for the Lexical Decision task.

### MRI Protocols and Preprocessing

MRI data were acquired using 3T Magnetom Prisma MR scanners. For functional scans, a multiband multi-echo sequence with a 32-channel head coil was used. Noise-cancelling headphones were used for presenting the auditory stimuli and reducing scanner noise and auditory responses were recorded with noise cancellation filtering. The following settings were used: TR = 1.50 s, TE1 = 13.4 ms, TE2 = 34.8 ms, TE3 = 56.2 ms, flip-angle = 75°, slice thickness = 2.5 mm, FOV = 210 mm, resolution = 2.5 x 2.5 x 2.5 mm^3^, slice matrix size = 84 x 84. For structural scans, MPRAGE T1-weighted anatomical scans at 1 mm isotropic resolution were acquired with a repetition time of 2000 msec, an echo time of 2.03 msec, a flip angle of 8°, and a field of view of 256 × 256 × 192 mm. All collected data (DICOM files) were converted to BIDS compatible format using bidscoiner (vers. 3.7.4 https://github.com/Donders-Institute/bidscoin). Functional data were further preprocessed using fmriprep (v22.1.1), with added function of normalizing to MNI152NLinAsym template with the voxel size of 2 mm^3^, and calculating the motion related independent components using AROMA function (97). All scans were smoothed with 4 mm kernel FWHM using SPM12 function (www.fil.ion.ucl.ac.uk/spm/software/spm12/).

### Laterality Index Calculation

To quantify hemispheric lateralization for each language task, laterality indices (LIs) were calculated for each participant based on participant-level T*-*maps. For each contrast image, voxels exceeding a T-value of 3.13 (corresponding to p = 0.001, two-tailed) were considered suprathreshold and included in the calculation. For each participant, LI was computed as

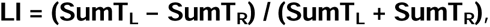

where SumT_L_ and SumT_R_ denote the sum of T-values across suprathreshold voxels in the left and right hemispheres, respectively. LI values ranged from −1 to +1, with negative values indicating right-lateralized activation and positive values indicating left-lateralized activation. At the group level, one-sample t-tests against zero were performed to assess whether lateralization was consistent across participants. Effect sizes were quantified using Cohen’s d, calculated as the mean LI divided by the standard deviation of LI across participants.

## Acknowledgments

We thank Xin Liu, Bob Kapteijns, Vera van’t Hoff, Marjolijn Dijkhuis, Jiska Koemans, Robert van Dongen, Milou Huismans, Frederieke Taheij, Levi Voeteé, Noëlle Bauhuis, Sascha Derks, Simone van Amerom, Janniek Wester, and Kyla McKonnell for their contributions to this work. These contributions included assistance with experimental setup, data collection, audio data transcription, and providing feedback on the manuscript.

This work was funded by the Dutch research council (Nederlandse Organisatie voor Wetenschappelijk Onderzoek) NWO Grant Language in Interaction, under Gravitation grant number 024.001.006

