## Supplementary Information for "The scaffolding of individual variability in language processing by domain-general neural networks"

**Lexical Decision Task - Stimuli:** The stimuli consisted of 80 existing Dutch nouns, 80 pseudowords and 20 quilts. The 80 existing words were chosen to differ in the word frequency on a continuum ranging from Zipf frequency values of 2.15 – 5.27, with a mean Zipf frequency of 3.71 (1). Additionally, the stimuli were selected based on several other psycholinguistic variables, including prevalence (> 0.959), word length (4 - 9 letters and 1 – 4 syllables), phonological neighborhood density (0 - 26), and phonological neighborhood frequency (0 – 3362.93). The 80 pseudowords for this task were generated by modifying the existing words using Wuggy software (2). Recordings of all the existing words and pseudowords were produced by one female speaker of Dutch. From the 80 existing words, 20 were selected (by sampling evenly from the full frequency range) to be transformed into “quilts”, to be used as baseline trials. Specifically, these quilts were created by dividing real word recordings into segments of 30 ms in length, which were then concatenated in a semi-random order. This semi-random concatenation of segments was performed such that it maximized the cross-correlation between the original and quilted waveforms. This quilting procedure was based on the “Sound Quilting Toolbox” (3), which essentially disrupted sound properties of audio stimuli on longer timescales, while approximately preserving them on shorter timescales. This means that quilted stimuli maintained their low-level auditory features (i.e., their speech-specific spectrotemporal structure), without any recognizable phonotactic or lexical features (**Figure S1**).

**Picture Naming Task - Stimuli:** The stimuli consisted of 80 pictures of common objects, 20 nameless objects and 20 scrambled pictures. The 80 pictures were taken from the Bank of Standardized Stimuli (4). All pictures were colored photographs located on a square white background, with equal dimensions and an equal digital resolution of 300 x 300 pixels. The objects were selected such that the name of the objects ranged in the word Zipf frequency values (range 2.06-4.84). Twenty of the 80 pictures were scrambled and used as baseline stimuli (5). Specifically, scrambled pictures were generated by dividing the intact pictures into a 20*20 grid of small squares (each 15*15 pixels), and by randomly rearranging the positions of these small squares. The resulting scrambled pictures were thus similar to the original pictures in terms of low-level visual features and complexity, but without any recognizable parts or objects. The twenty original pictures used for scrambling were selected by sampling evenly from the visual complexity distributions in all pictures, whilst ensuring that the selection had sufficient variation in terms of color schemes. Twenty nameless objects were selected from other studies that used such objects in their studies (6, 7) to be included as a baseline condition. This condition required the visual processing of picture stimuli and generating a verbal response (“Niks”) without engaging in accessing, retrieving, or preparing lexical representations for production (**Figure S2**).

**Sentence Comprehension Task - Stimuli:** 80 sentences were selected from a published data set Mother Of Unification Studies (MOUS) created by Schoffelen et al. (8) and were used in a study by Uddén et al. (9). All sentences varied between 9 and 15 words in length, and they were relatively unconstrained in terms of syntactical structure. Forty-six of the 80 selected sentences contained a relative clause (58%), whereas 34 did not (42%). Word frequencies for each word in each sentence were retrieved from SUBTLEX-NL (2), and converted to the Zipf scale (1). For each sentence, we calculated the average Zipf frequency, as well as the minimum Zipf of all words within each sentence. For the word list condition, 140 words were randomly selected from the above list of sentences. These words were distributed amongst 20-word lists with seven words in each list. The mean Zipf frequency values of the words per list and the minimum Zipf frequency value within the list was similar to that of the sentence condition on average. In 20% of the cases (16 trials for sentence condition, and 4 trials for word list condition), a question followed on the screen, which focused on the content of the sentence, or whether the word was mentioned in the preceding sentence / list. (e.g. “Was a profession mentioned?”, “Was the word ‘barkeeper’ mentioned?”). All sentences were additionally characterized in terms of syntactic complexity, as well as word count. The syntactic complexity of each sentence was assessed using a left-branching complexity measure from a dependency structure (9, 10). We used a left-branching unification score, which counts the number of left-branching dependents that are closed at each word in a dependency structure (for example, in the sentence “the customer left”, one dependency relation (determiner to noun) is closed at “customer”, and one relation (subject to verb) is closed at “left”). The scores were summed across all words in a sentence, reaching a total unification score for each sentence (**Figure S3**).

**Sentence Production Task - Stimuli:** The stimuli were adapted from the set of photographs used in the studies by Menenti et al. (11, 12). The stimulus set included 30 images depicting a single person shaded in yellow performing an action, 30 images showing two people (a man and a woman) engaged in an action with one shaded in yellow and the other in blue, and 30 additional images featuring the same pair performing actions with the color assignments swapped between the man and the woman. In addition, 30 intransitive, 30 active and 30 passive sentences, and two phrases spoken by a native Dutch young male were used. In total there were 30 active trials, 30 passive trials, 30 intransitive trials and 24 phrase trials (baseline) (**Figure S4**).

**Figures**

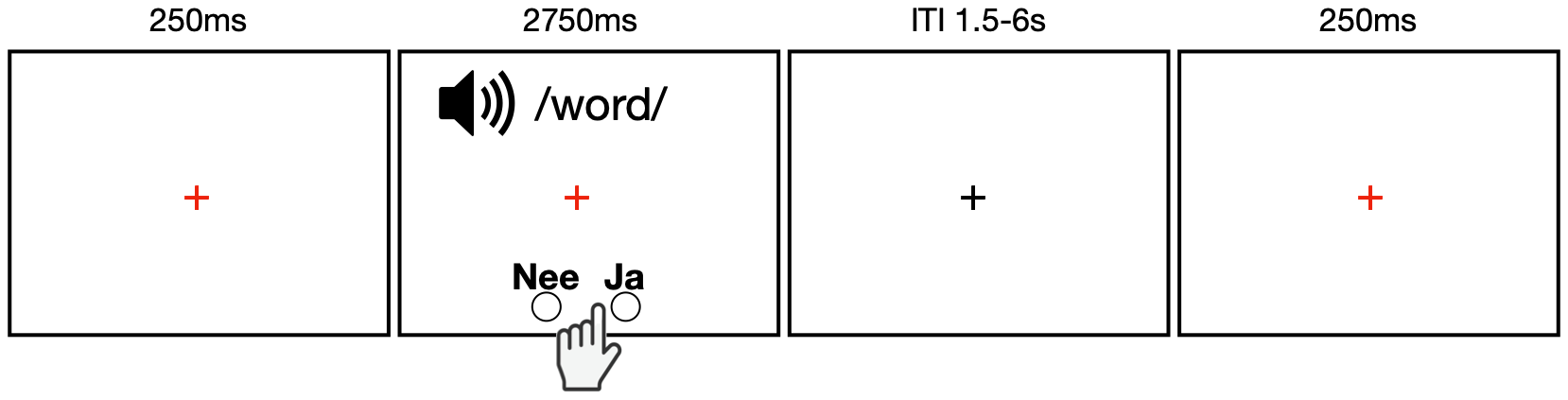

Figure S1. Example trial structure for the auditory lexical decision task

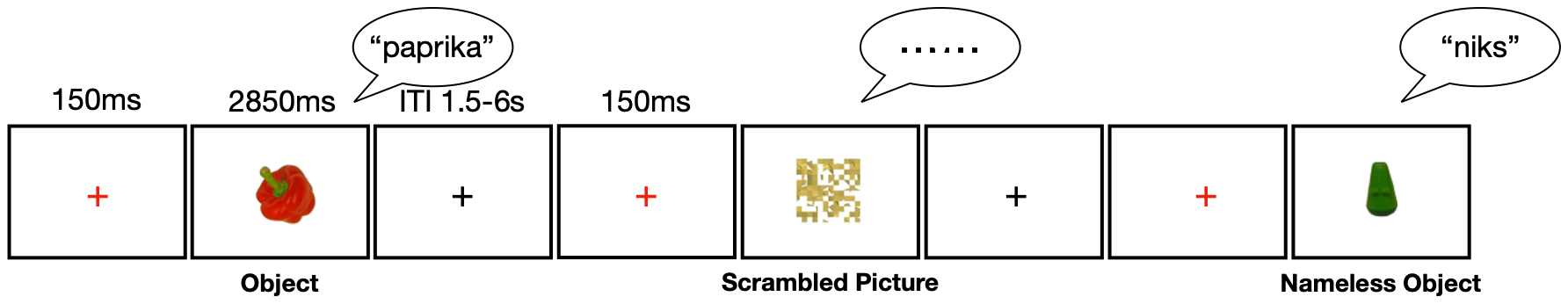

Figure S2. Example trial structure for the picture naming task

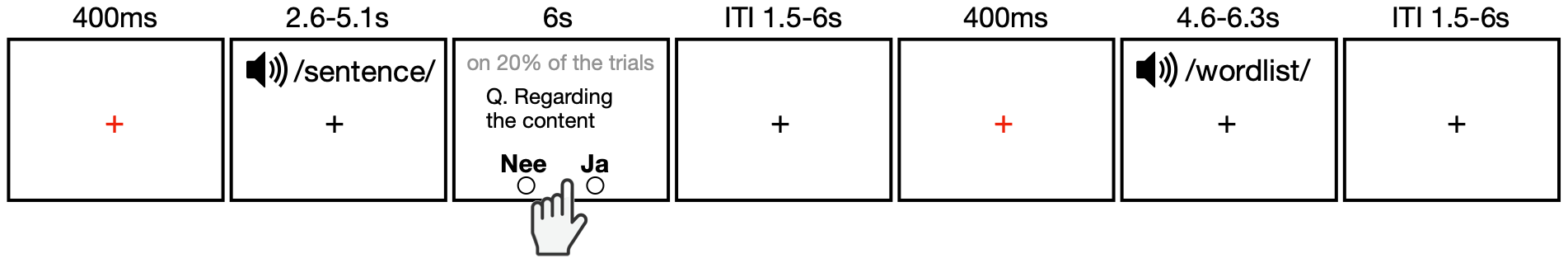

Figure S3. Example trial structure for the sentence comprehension task

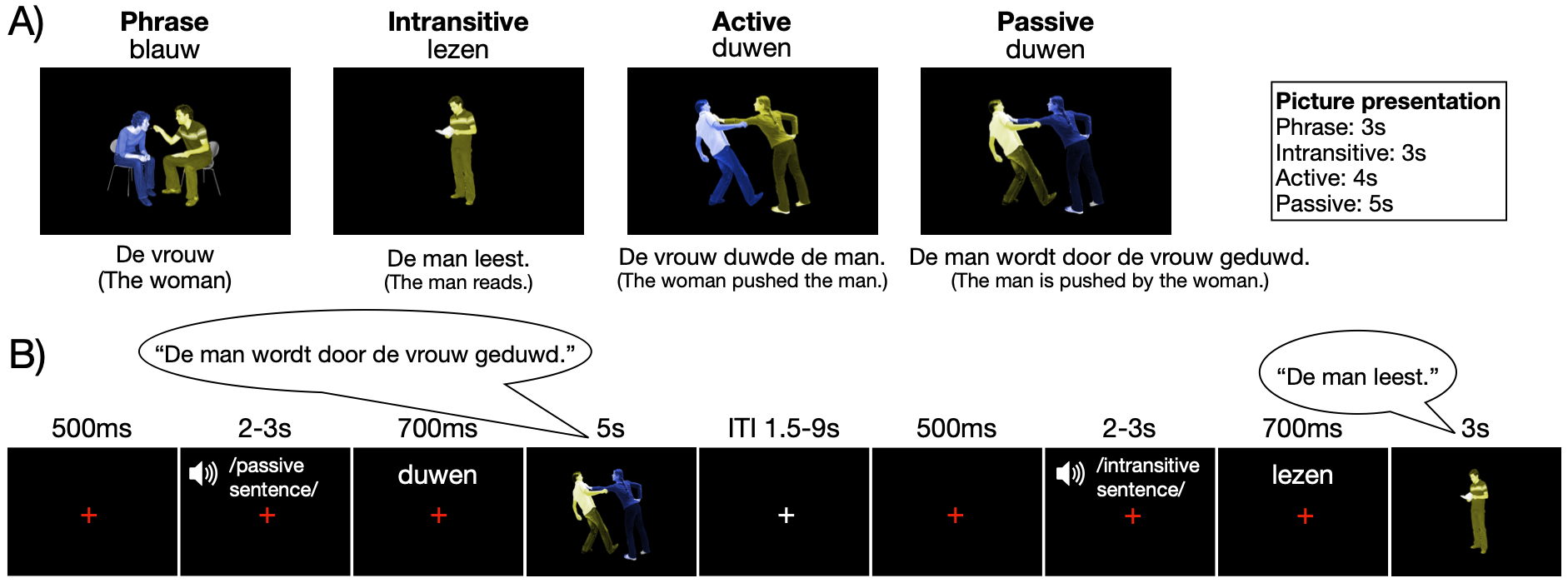

**Figure S4.** Example **A.** stimuli and **B.** trial structure for the sentence production task

Tables

**Table S1.** Behavioral tests included in the battery used to assess the four cognitive skills as in Hintz et al. (13)

| Linguistic Knowledge | Processing Speed | Working Memory | Non-verbal Reasoning |
| --- | --- | --- | --- |
| Peabody picture vocabulary  Antonym production  Idiom recognition  Spelling  Author recognition  Prescriptive grammar | Auditory simple reaction time  Auditory choice reaction time  Letter comparison speed  Visual simple reaction time  Visual choice reaction time | Digital forward span  Digital backward span  Corsi forward block clicking  Corsi backward block clicking | Raven’s advanced progressive matrices |

Table S2. Behavioral accuracy scores of the fMRI language tasks averaged across participants

| **Language Task** | **Condition** | **Mean Accuracy Score (%)** |
| --- | --- | --- |
| **Lexical Decision** | Word | 94 |
|  | Pseudoword | 91 |
|  | Quilt | 100 |
| **Picture Naming** | Object | 94 |
|  | Nameless Object | 98 |
|  | Scrambled Picture | 100 |
| **Sentence Comprehension** | Sentence | 96 |
|  | Wordlist | 94 |
| **Sentence Production** | Active | 95 |
|  | Passive | 94 |
|  | Intransitive | 96 |
|  | Phrase | 96 |

**Table S3.** Significant fMRI group-level activation clusters across different language task contrasts and modulators (thresholded at a voxel-wise level of p < 0.001, with cluster-level Family-Wise Error correction at p < 0.05). Peak coordinates are reported in MNI space, and anatomical labels were assigned using the Automated Anatomical Labeling (AAL) atlas implemented in SPM12.

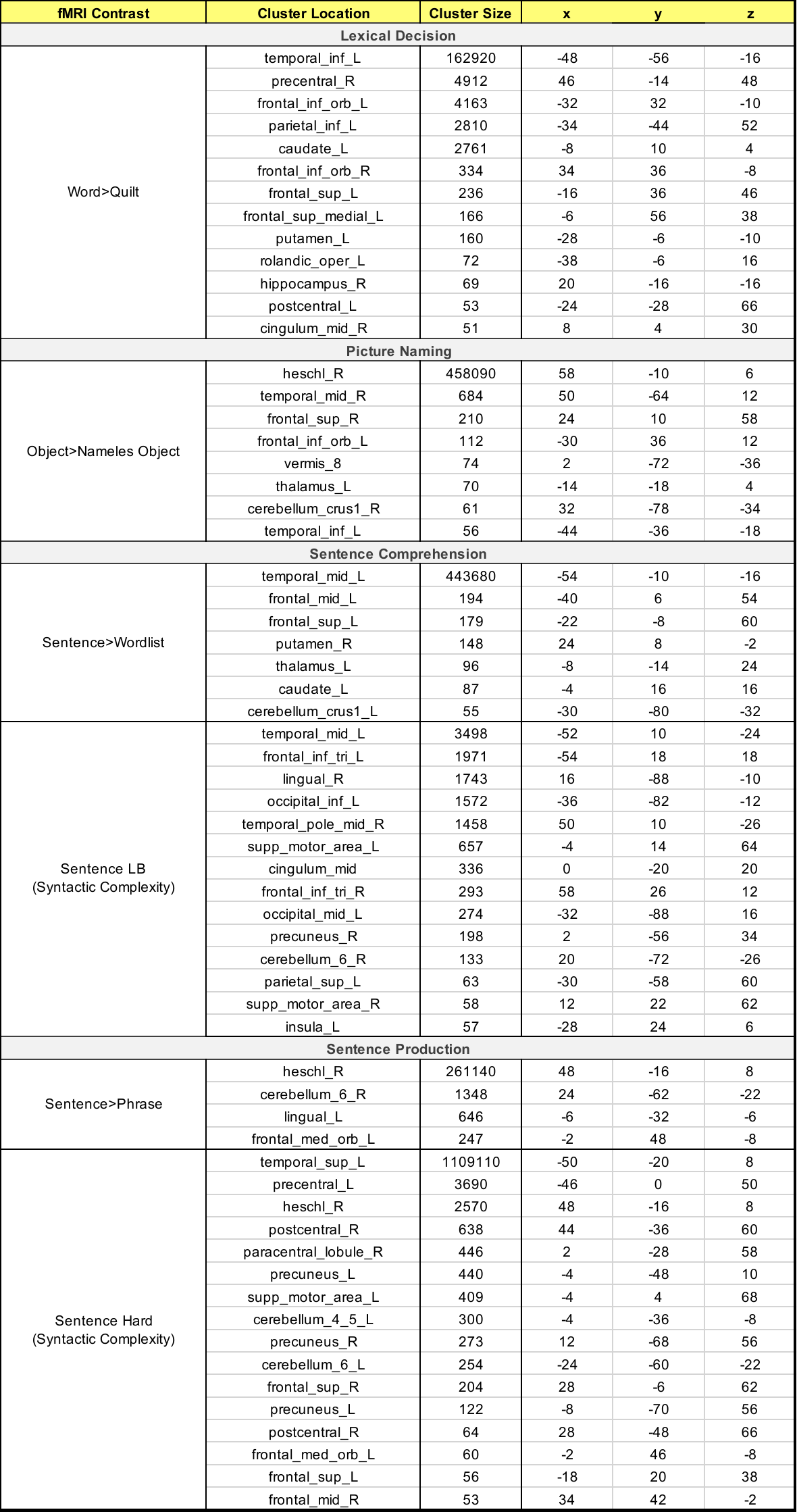

**Table S4.** ‘**Individual variability clusters’ where fMRI activation covaried with cognitive skills.** The **Contrast/Modulation** column specifies the experimental fMRI condition contrast or parametric modulator used for each language task. **P-FWE** denotes the cluster-level Family-Wise Error-corrected significance level. **Size** denotes cluster size in number of voxels. **x, y, z** columns indicate the MNI coordinates of the peak voxel within each cluster. **Overlap** quantifies the percentage of spatial overlap between the cluster and the group-level common activation map. SMA: Supplementary motor area, RO: Rolandic Operculum, IFG op.: Inferior frontal gyrus opercularis, MTG: Middle temporal gyrus, STG: Superior temporal gyrus, MFG: Middle frontal gyrus, SMG: Supramarginal gyrus, AG: Angular gyrus, IPL: Inferior parietal lobule, med. SFC: Medial superior frontal cortex, ACC: Anterior cingulate cortex, sup. Occ: superior occipital gyrus, R: right, L: left

| Cluster | Contrast/ Modulation | Cognitive Skill | Correlation | Region | P-FWE | Size (voxels) | x | y | z | Overlap |
| --- | --- | --- | --- | --- | --- | --- | --- | --- | --- | --- |
| **Lexical Decision** | | | | | | | | | | |
| 1 | Word>Quilt | Linguistic Knowledge | Negative | SMA (R) | 0 | 151 | 4 | -4 | 68 | 22% |
| 2 | Word>Quilt | Linguistic Knowledge | Negative | RO (R) | 0.041 | 53 | 62 | 8 | 12 | 0 |
| 3 | Word>Quilt | Processing Speed | Positive | Cerebellum Crus I (L) | 0.007 | 74 | -38 | -50 | -30 | 26% |
| 4 | Word>Quilt | Processing Speed | Positive | Cerebellum VI (R) | 0.049 | 51 | 22 | -58 | -20 | 96% |
| 5 | Word>Quilt | Non-verbal Reasoning | Negative | IFG op. (L) | 0.044 | 52 | -58 | 14 | 12 | 23% |
| **Picture Naming** | | | | | | | | | | |
| 6 | Object>Nameless Object | Processing Speed | Negative | MTG (L) | 0.032 | 58 | -48 | -56 | 4 | 0 |
| 7 | Object>Nameless Object | Working Memory | Negative | Precuneus (L) | 0.053 | 52 | 0 | -66 | 42 | 0 |
| **Sentence Comprehension** | | | | | | | | | | |
| 8 | Sentence>Wordlist | Linguistic Knowledge | Negative | IPL (L) | 0 | 131 | -40 | -48 | 38 | 0 |
| 9 | Sentence>Wordlist | Linguistic Knowledge | Negative | Cerebellum Crus I (R) | 0.012 | 71 | 40 | -62 | -30 | 0 |
| 10 | Sentence>Wordlist | Linguistic Knowledge | Negative | Insula (R) | 0.031 | 59 | 32 | 22 | 12 | 0 |
| 11 | Sentence>Wordlist | Linguistic Knowledge | Negative | MFG (L) | 0.013 | 70 | -50 | 12 | 38 | 0 |
| 12 | Sentence>Wordlist | Processing Speed | Negative | MTG (L) | 0.016 | 67 | -48 | -52 | 16 | 73% |
| 13 | Sentence>Wordlist | Processing Speed | Negative | STG (R) | 0.031 | 59 | 52 | -4 | -4 | 76% |
| 14 | Syntactic Complexity | Non-verbal Reasoning | Positive | Cerebellum Crus I (L) | 0.006 | 73 | -18 | -56 | 32 | 0 |
| 15 | Syntactic Complexity | Processing Speed | Negative | MFG (L) | 0.04 | 51 | -28 | -60 | 40 | 0 |
| Sentence Production | | | | | | | | | | |
| 16 | Sentence>Phrase | Working Memory | Positive | Calcarine (L) | 0.004 | 88 | -16 | -80 | 8 | 11% |
| 17 | Sentence>Phrase | Working Memory | Negative | Precuneus (L) | 0.002 | 96 | 0 | -68 | 44 | 0 |
| 18 | Sentence>Phrase | Non-verbal Reasoning | Negative | MFG (L) | 0.026 | 63 | -36 | 46 | 0 | 0 |
| 19 | Syntactic Complexity | Linguistic Knowledge | Positive | med. SFC (L) | 0.011 | 71 | -8 | 54 | 14 | 0 |
| 20 | Syntactic Complexity | Linguistic Knowledge | Positive | ACC (L) | 0 | 176 | -4 | 36 | 0 | 2% |
| 21 | Syntactic Complexity | Linguistic Knowledge | Positive | Calcarine (R) | 0.006 | 79 | 12 | -82 | 16 | 11% |
| 22 | Syntactic Complexity | Linguistic Knowledge | Positive | sup. Occ (L) | 0 | 119 | -16 | -86 | 32 | 0 |
| 23 | Syntactic Complexity | Linguistic Knowledge | Positive | IPL (L) | 0.046 | 53 | -58 | -44 | 40 | 0 |
| 24 | Syntactic Complexity | Processing Speed | Negative | SMG (L) | 0.001 | 112 | -58 | -34 | 32 | 0 |
| 25 | Syntactic Complexity | Non-verbal Reasoning | Negative | ACC (R) | 0.006 | 79 | 4 | 46 | 16 | 0 |
